# MAHLER: Integrating Metadynamics and Inverse Folding to Predict Antibody-Antigen Kinetics

**DOI:** 10.64898/2026.06.10.731383

**Authors:** Da Teng, Mary Pitman, Punit K. Jha, Amogh Sood, Dominic Rufa, Kevin Ryczko, Andrea Bortolato, Pratyush Tiwary

## Abstract

Binding kinetics are crucial for antibody function, shaping pharmacokinetics and *in vivo* efficacy beyond what equilibrium affinity captures. We present “Metadynamics-Anchored Hybrid Learning for Engineering off-Rates (MAHLER)”, a fully open-source machine learning/physics hybrid method that predicts relative antibody-antigen residence times at scale. Incorporating inverse-folding models into molecular dynamics simulations, MAHLER shows first-in-class screening-grade accuracy in calculating relative antibody-antigen dissociation kinetics across a family of point mutants. After initial antigen-specific setup, each prediction takes only 4 minutes on a single NVIDIA A100 GPU, compared to days even with already enhanced molecular dynamics simulations. This provides practical kinetics-aware complement to current computational design approaches that focus primarily on binding affinity for antibody-antigen complexes.

## Introduction

AI-assisted computational antibody design is maturing rapidly, with state-of-the-art models now achieving double-digit success rates in *in vitro* binding affinity validation^1–3^. Despite this progress, the kinetic profile of antigen-antibody (AgAb) binding remains a critical blind spot in computational design. Binding affinity alone is often insufficient to predict *in vivo* therapeutic outcomes^4, 5^. However, the dissociation rate (*k*_off_), or the inverse of the residence time (*τ*), dictates the duration of biological interaction, thereby governing therapeutic efficacy and pharmacokinetics^6^. Additionally, in diagnostics, longer residence time is the primary driver of sensitivity in ultrasensitive immunoassays^7^.

Kinetics-aware AI design of antibodies, while desirable, remains difficult for numerous reasons^5^. First, experimental data for kinetics is scarce and often noisy due to external factors like non-specific binding, which limits the applicability of purely supervised machine learning techniques^8, 9^. Second, current structure-based models lack the sensitivity needed to reliably predict the subtle effects of single-point mutations^10^, which are crucial in antibody engineering. Some differences are also not reflected in the static bound structure, but rather manifest in the physical interactions that occur during the dissociation process^4^. While physics-based sampling such as molecular dynamics (MD) can theoretically capture these nuances, it suffers from severe sampling bottlenecks. While standard unbiased MD is typically limited at its very best to several hundred microseconds, the dissociation of therapeutically relevant antibodies often occurs on the seconds timescale, placing it far beyond the practical reach of these simulations^11^.

Infrequent metadynamics (IfMetaD) offers a theoretical framework within the metadynamics family of enhanced sampling methods to bridge this gap, predicting residence times primarily for protein-ligand systems^12–15^, and more recently, protein-protein complexes^16, 17^. Despite its potential, the low computation throughput of IfMetaD hinders its broader application in prospective antibody design. Most importantly, prospective design requires screening a large number of candidate mutants. While automated protocols have been recently developed for calculating absolute protein-small molecule residence times^14, 15^, IfMetaD remains prohibitively expensive for high-throughput screening of antibody-antigen kinetics. Meanwhile, IfMetaD relies on high-quality collective variables (CVs) to accurately describe the dissociation process. Identifying CVs that capture relevant slow degrees of freedom often requires extensive trial-and-error simulation, which has been shown feasible for protein-small molecule systems^14, 15, 18^ but remains exceptionally challenging for macromolecular protein-protein interfaces^19–21^.

Here, we describe a first-in-class protocol (Figure 1a), called Metadynamics-Anchored Hybrid Learning for Engineering off-Rates (MAHLER), which integrates AlphaFold2-RAVE (af2rave)^22, 23^, IfMetaD^12^, and inverse folding models (IFM)^24^ to efficiently sample the dissociation of antibody-antigen complexes. For a given wild-type (WT) complex, af2rave generates a reduced-dimensional latent space that encodes slowly interconverting conformations possibly relevant to the dissociation process (Figure S4). Applying IfMetaD within this optimized latent space, with the learned latent variables serving as the collective variables (CVs) for biasing, allows us to accelerate dissociation events and accurately estimate residence times for the WT complex (Figure 1c). By incorporating the IFM framework, MAHLER eliminates the need for repeated IfMetaD simulations on each individual mutant by reweighting the single WT IfMetaD trajectory with much faster inverse folding model inferences, directly addressing the aforementioned challenges. Detailed simulation parameters and CV generation protocols are provided in the Methods section.

**Figure 1.**
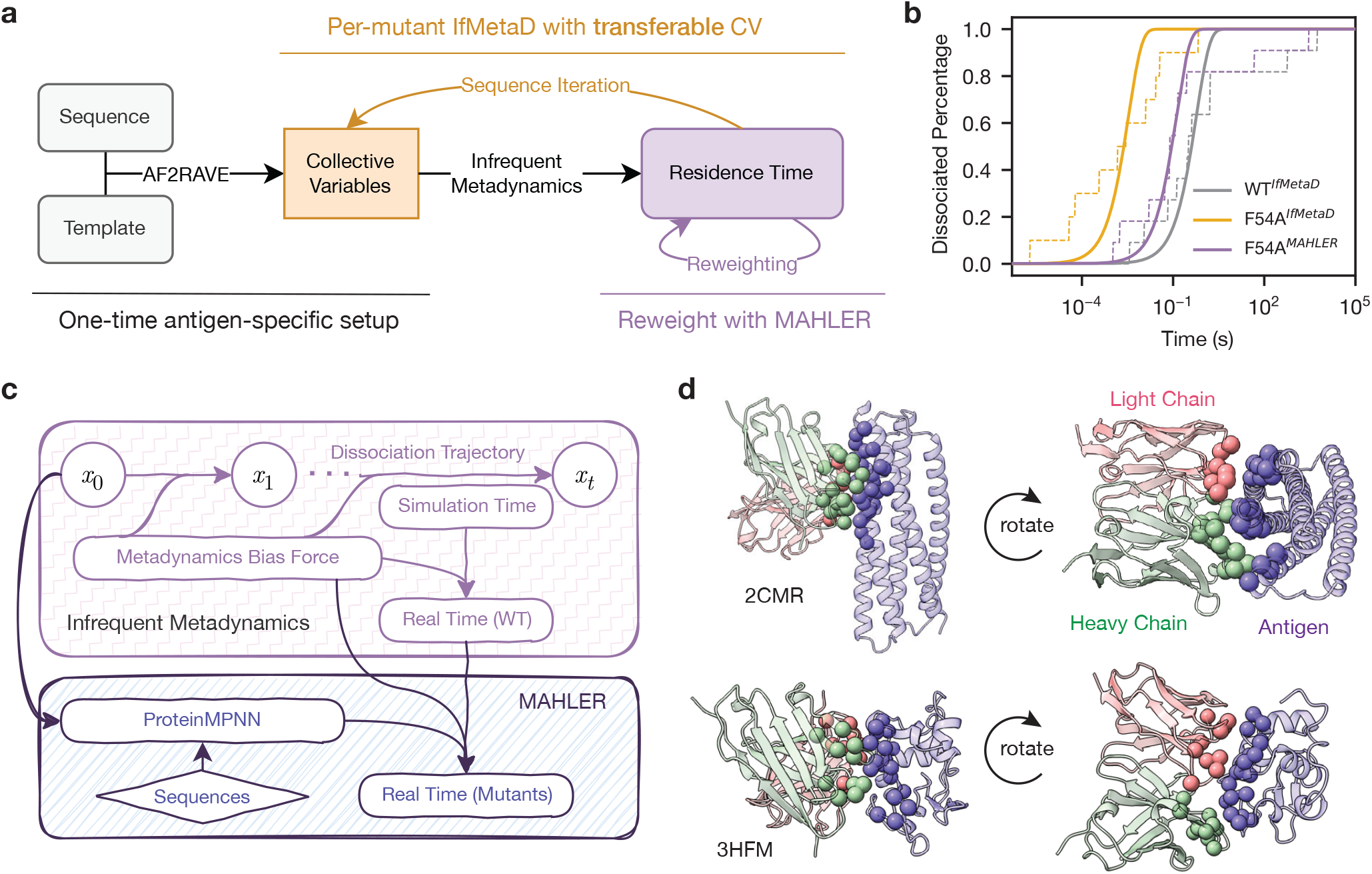
Workflow of MAHLER and its validation systems. (a) Overview of the computational workflow described in this paper. The orange and purple color scheme is used consistently throughout the manuscript to distinguish between methods. An example is shown in panel (b), where the gray curve corresponds to an IfMetaD simulation of the wild-type (WT) sequence of 2CMR, serving as a one-time, antigen-specific setup. The purple curve is a mutant F54A reweighted with MAHLER from the WT trajectories, and the orange curve is derived from per-mutant IfMetaD using CVs learned from WT sampling. (c) Illustration of regular IfMetaD and MAHLER. A trajectory of molecular coordinates *x*_0_, · · ·, *x*_*t*_ is sampled from IfMetaD, and the metadynamics bias force along the trajectory is used to compute the acceleration factor *α*, which connects the simulation time to the real dissociation time. MAHLER takes this trajectory and a mutant sequence to compute a log-likelihood score with ProteinMPNN. These weights are integrated with the bias force to compute the acceleration factor for the mutants, yielding the mutant-specific dissociation time. (d) Side and top views of the two validation systems. Interface residues are marked with spheres.

## Results

### Metadynamics-Anchored Hybrid Learning for Engineering off-Rates (MAHLER)

While similar approaches have proven successful in small ligand-protein residence time predictions, they are not yet applicable to antibody design problems^15^. In antibody design, the combinatorial space of sequence hypotheses that must be tested is vast, demanding highly efficient approaches for rapid, high-throughput screening. To bridge this gap, we developed MAHLER to circumvent the prohibitive computational cost of repeated, per-mutant IfMetaD calculations by efficiently reweighting a single WT trajectory using sequence likelihoods derived from an inverse folding model (IFM). As derived in the Methods section 5 building on Ref. 17, Bayes’ theorem allows us to relate the conditional probabilities of WT and mutant sequences to a particular conformation, meaning that sampling the target mutant ensemble can be achieved by applying ProteinMPNN scores as statistical weights to the pre-computed WT trajectories (Equation (5)).

We validated MAHLER on three sets of point mutations across two AgAb systems (Figure 1d) to rigorously evaluate its predictive capabilities and establish its operational boundaries. The two systems selected feature antibodies neutralizing (a) the gp41 protein inner-core mimetic of human HIV-1 (`pdb_00002cmr`), and (b) the hen eggwhite lysozyme (HEL, `pdb_00003hfm`). For the gp41 system, both the experimental structure and kinetics measurements were reported in reference 25. The kinetics data were derived from an alanine scan that mutated 12 residues of the antibody to alanine to probe the importance of interfacial residues. In the HEL system, structures^26^ and kinetic measurements^27–29^ were reported separately. The kinetics data we selected comprise two sets of mutations where two important residues on the antigen were systematically mutated to different amino acids.

For each validation system, we computed or compiled three sets of data: (1) the experimental *k*_off_ from the literature, (2) per-mutant IfMetaD calculations utilizing the transferable CVs learned by af2rave, and (3) the reweighted time derived from the WT IfMetaD calculation using MAHLER. By analyzing the correlation among these metrics, we systematically assess the fidelity of our reweighting scheme against brute force simulations and ground-truth experiments. In our brute force calculations, we learned the CV from conformational sampling of the WT AgAb complex, then transfer them to mutants. Since the CVs learned by af2rave used in these simulations are linear combinations of distances between C_*α*_ atoms of interface residues, they can be seamlessly transferred across mutants.

### MAHLER predictions agree with per-mutant calculations

Our calculations show that the relative *k*_off_ predicted by MAHLER strongly correlates with per-mutant calculations (Figure 2a-c), demonstrating its viability as an efficient replacement for expensive, repetitive IfMetaD calculations for antibody-antigen kinetics. This validates our core hypothesis: evaluating structure-conditioned sequence likelihoods from an IFM within our trajectory reweighting framework provides sufficient information to reweight a single sampling trajectory and accurately infer kinetic properties of unseen mutants. The Spearman coefficient *r*_*s*_ (Table 1) quantifies this success, revealing robust correlations (~ 0.6) for the D101 mutants of 3HFM (Figure 2b) and 2CMR (Figure 2c), and a moderate but statistically meaningful rank-ordering capacity for the R21 mutants of 3HFM (Figure 2a). This level of accuracy in predicting relative unbinding kinetics of antibody-antigen systems is comparable with the state-of-the-art accuracy recently demonstrated in protein-small molecule absolute unbinding kinetics prediction.^14^ Notably, within the 3HFM dataset, the significant outliers (such as the R21E and D101R mutants) arise consistently from charge flipping (Figure 2a–b), a persistently difficult problem in other methods, including free energy perturbation (FEP)^30, 31^.

**Table 1.** Correlations between different measurements. In the three systems we computed the Spearman correlation (*r*_*s*_) and Pearson correlation (*r*_*p*_) between 3 different pairs of measurements from MAHLER, IfMetaD, and experiment.

| Pair of measurements |  | 3HFM (R21) |  | 3HFM (D101) |  | 2CMR (Ala-scan) |  |
| --- | --- | --- | --- | --- | --- | --- | --- |
| | | $r_s$ | $r_p$ | $r_s$ | $r_p$ | $r_s$ | $r_p$ |
| MAHLER | IfMetaD | 0.381 | 0.253 | 0.600 | 0.573 | 0.643 | 0.547 |
| MAHLER | Experiment | −0.299 | −0.508 | 0.657 | 0.316 | 0.490 | 0.483 |
| IfMetaD | Experiment | 0.527 | 0.368 | 0.086 | 0.220 | 0.685 | 0.629 |

**Figure 2.**
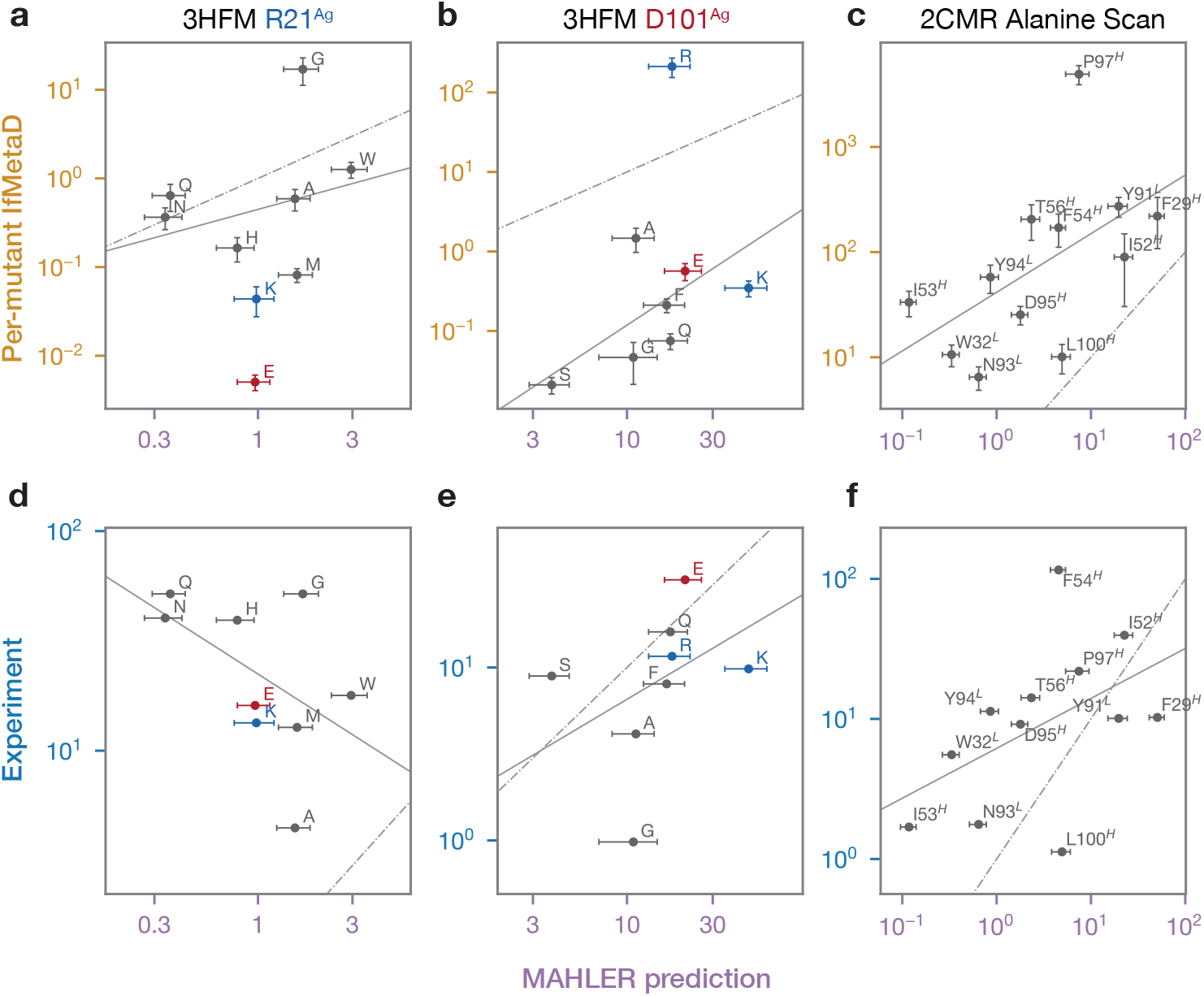
Performance of predictive methods. Each panel plots the relative *k*_off_ of a set of mutants, and a solid line for the best linear fit (without charge flipping), as well as a dashed line for *y* = *x* that is the perfect prediction. The error bars are from bootstrapping the individual samples 1000 times as detailed in Methods section. (a-c) compares MAHLER’s reweighting results with per-mutant IfMetaD calculation. (d-f) compares MAHLER predictions with experimental data.

### Benchmarking MAHLER against experimental kinetics identifies sampling limits

However, when benchmarked directly against experiments, MAHLER exhibits system-dependent levels of agreement (Figure 2d–f, Table 1). For 2CMR, the correlation achieves screening-grade accuracy capable of informing antibody design, with both *r*_*s*_ and Pearson correlation *r*_*p*_ consistently high at ~ 0.5. In contrast, performance on the 3HFM system drops significantly. The model fails to capture the trend for R21 mutants, while for D101 mutants, it successfully predicts the ranking (*r*_*s*_) but struggles to reproduce the numerical correlation (*r*_*p*_).

Fundamentally, MAHLER integrates two primary inputs (Figure 3a): sampling trajectories from IfMetaD and scoring weights from the IFM. For predictions to achieve fidelity with experimental data, both components must be highly reliable. Supporting this requirement, evaluation of the two 3HFM validation sets reveals that where MAHLER’s accuracy degrades, the underlying baseline per-mutant IfMetaD calculations similarly exhibit a poor correlation with experimental benchmarks (Figure 3c-d). Given the robust correlation observed between MAHLER and the reference per-mutant calculations (Figure 2a–c), the IFM weights are clearly excluded as the primary accuracy bottleneck, allowing us to turn our focus entirely to the underlying sampling quality. MAHLER succeeds on the antigen-antibody interaction in the 2CMR system, which is predominantly hydrophobic, while struggling on the 3HFM interface, which features more than ten pairs of polar interactions. This chemical complexity makes it very difficult to fully describe the dissociation of the 3HFM complex using only two CVs. The dissociation pathway of 3HFM likely involves a combinatorially large number of intermediate states characterized by different states of hydrogen bonds and other interactions forming/breaking. Consequently, these states can be difficult to describe adequately with low-dimensional CVs, a limitation signified by the weak correlation between the baseline IfMetaD calculations and experiments.

**Figure 3.**
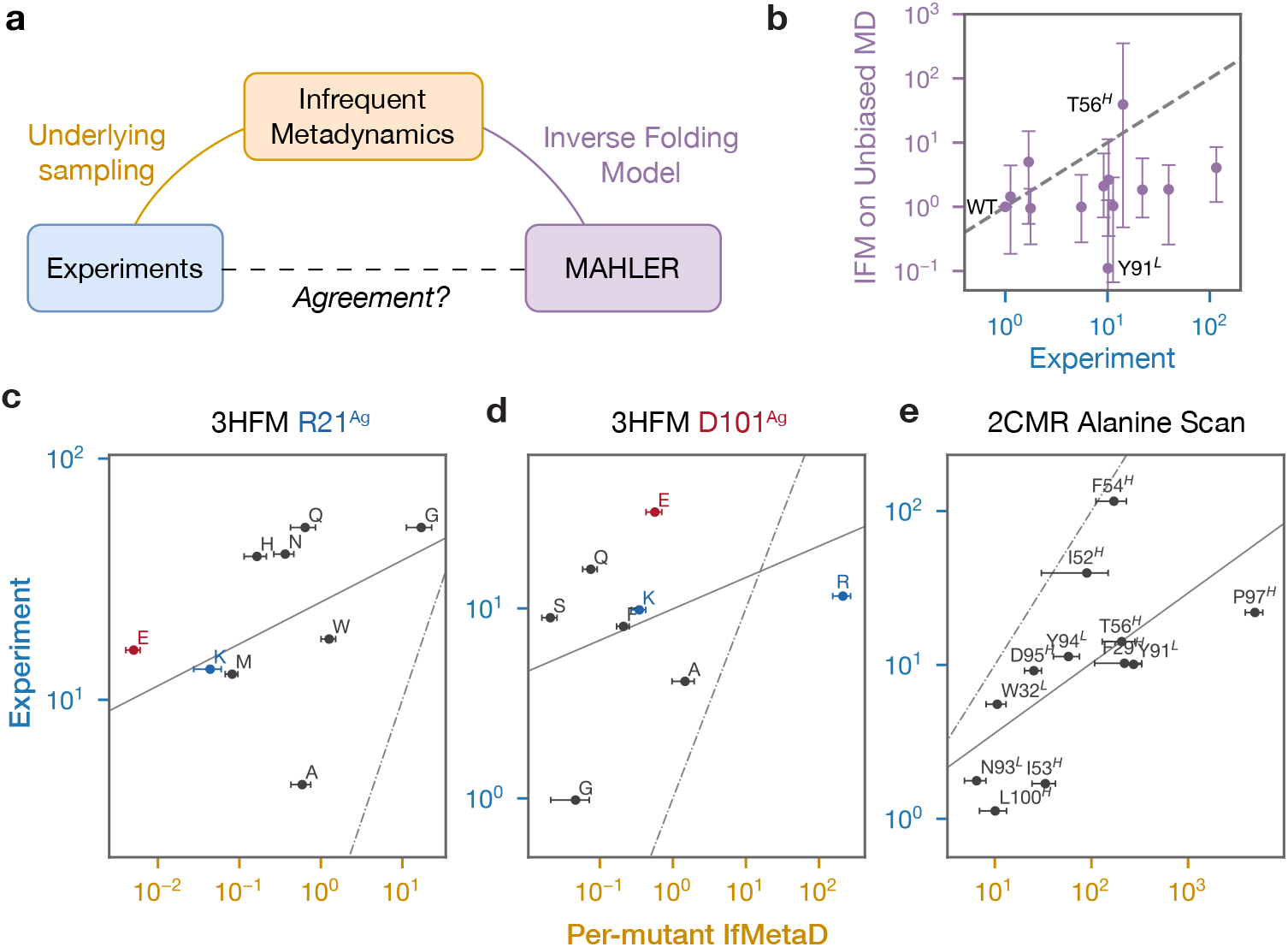
Dissecting the assumptions of MAHLER and its accuracy. (a) MAHLER’s accuracy relies on two key factors: the quality of the underlying sampling of IfMetaD and the accuracy of the Inverse Folding Models. (b) The average IFM score from an unbiased trajectory from the experimental structure of 2CMR, plotted against the experimental *k*_off_ value. Error bar is from mixing three replicas of sampling and bootstrapping 1000 times with replacement. (c-e) Comparing MAHLER’s prediction directly with experimental measurements in three validation systems. Solid line is the linear fit and the dashed line is *y* = *x* which is the perfect agreement.

To definitively prove that capturing the explicit physical unbinding trajectory is what drives this predictive capacity, rather than simple static sequence matching, we scored unbiased MD simulations initiated from the experimental 2CMR complex structure. Without observing the dissociation process, the mean ProteinMPNN scores alone fail entirely to correlate with the experimental *k*_off_ (Figure 3b). This confirms that IfMetaD’s conformational sampling of the dissociation pathway is a strict prerequisite for predictive success. This failure mirrors recent findings in protein folding, where unbiased MD was similarly shown to yield no benefit for IFM-based stability predictions^32^. Ultimately, these results demonstrate that mere equilibrium fluctuations are insufficient and samples of the physical dissociation process is indispensable for accurate kinetic predictions.

However, this requirement does not necessitate flawless sampling to achieve engineering utility. Instead, MAHLER provides a highly practical and scalable compromise by utilizing af2rave to rapidly extract transferable CVs from bound-state data. Because MAHLER’s reweighting demonstrates system-independent agreement with explicit per-mutant calculations (Figure 2a–c), the framework establishes a self-consistent method for validation. In a prospective library screen, one can cross-check the high-throughput reweighted profiles by running explicit IfMetaD simulations on a small subset of variants. If the explicit simulation times align with the reweighted predictions, it provides immediate statistical confirmation that the underlying af2rave CV has adequately resolved the slow degrees of freedom governing that specific sequence space. Conversely, whenever highly disruptive changes such as localized charge-flipping mutations introduce structural uncertainty, this targeted physical validation can be selectively deployed as an internal diagnostic check to safeguard the accuracy of the screen.

### Performance of reweighting is hundreds of times faster than per-mutant IfMetaD

MAHLER allows us to achieve breakthrough efficiency in sequence screening. Specifically for the 2CMR system, this reweighting method is 760 times cheaper for each mutant than repeated IfMetaD simulations. Such a drastic reduction in computational cost makes it especially useful for prospective designs. On average, a data point from trajectory reweighting costs only 0.064 GPU hours on an NVIDIA A100, whereas the explicit calculations with IfMetaD take 48.7 GPU hours each. After an initial setup, the reweighting calculation becomes around 20 times faster in wall clock time than wet-lab experiments (96-minute half-life for 2CMR WT^25^). At the scale of a library screen of ~ 10^5^ sequences, the time saving is around 4 × 10^6^ GPU hours and millions of dollars (Figure 4).

**Figure 4.**
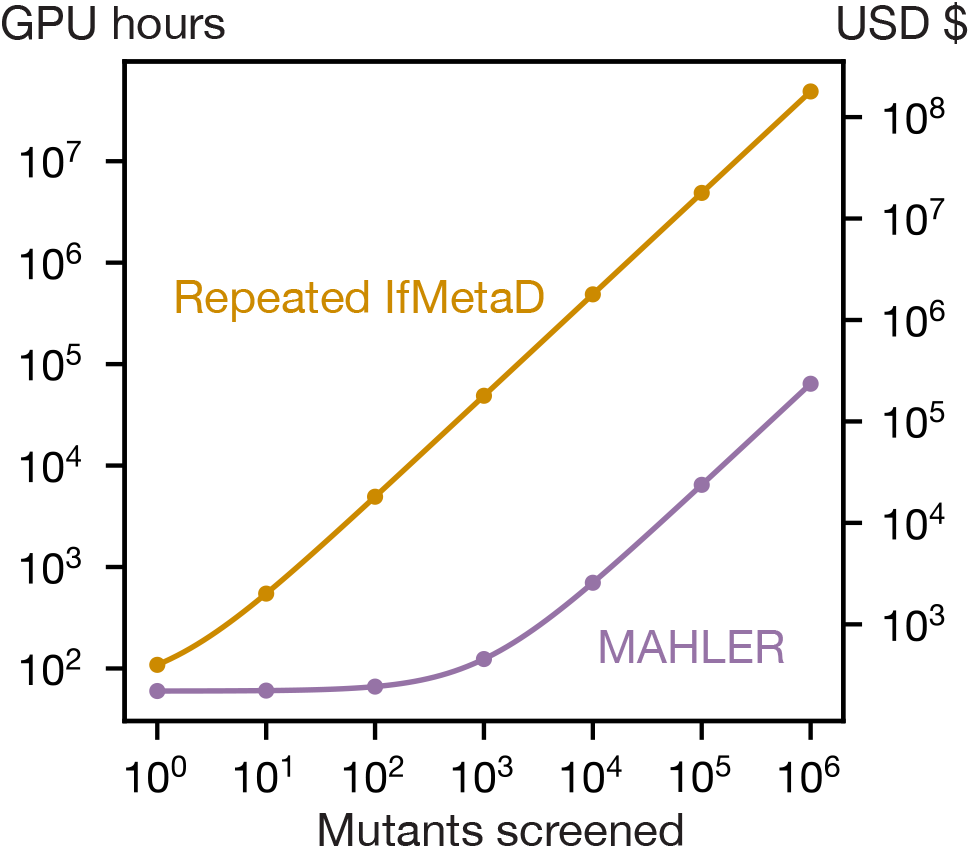
Computation cost of MAHLER compared with brute-force screening. The cost is estimated on NVIDIA A100 cost of $3.67/hour at Google Cloud Platform `a2-highgpu-1g`.

## Discussion

Predicting kinetics is intrinsically a formidable challenge, yet it remains extremely important for rational antibody design. Inferring kinetic rates from a static experimental structure is analogous to reconstructing an entire temporal trajectory from a single static observation. To address this gap, we introduce MAHLER, a framework that combines AF2RAVE for CV learning with a reweighting scheme utilizing protein IFMs. We demonstrate that an IFM coupled with enhanced sampling can serve as a highly efficient replacement for computationally expensive, per-mutant IfMetaD calculations. A core contribution of this work is demonstrating that rigorous enhanced sampling is a prerequisite for predicting kinetics, even when deploying sophisticated machine learning models trained on structural data, as scoring static equilibrium fluctuations yields no correlation with experimental *k*_off_ (Figure 3b). Furthermore, the baseline sampling quality dictates whether MAHLER achieves quantitative agreement with experimental benchmarks, as demonstrated by the screening-grade rank-ordering achieved for 2CMR (*r*_*s*_ ~ 0.5) compared to the challenges encountered at the polar 3HFM interface.

To understand the system-dependent performance of MAHLER, we need to examine the contrasting biochemical profiles of our validation targets. The framework delivers screening-grade accuracy for the 2CMR system, where the experimental alanine scan probes a predominantly rigid, hydrophobic interface. In contrast, the 3HFM interface is governed by a cooperative network of over ten polar interactions, systematically mutated to introduce complex hydrogen-bonding variations and charge-flipping events. Conceptually, trajectory reweighting requires that the structural conformations explored by the wild-type ensemble remain highly representative of those preferred by the mutants. Because the hydrophobic substitutions in 2CMR maintain structural homology with the wild-type unbinding pathway, the reference trajectory provides an excellent foundation for kinetic inference. However, disrupting the intricate polar networks or flipping localized charges in 3HFM forces the mutant complexes to seek entirely alternative dissociation pathways, visiting transition states and intermediates that are fundamentally absent from the wild-type simulation. Crucially, the strong agreement observed between MAHLER and explicit per-mutant IfMetaD calculations across both systems confirms that the inverse folding model’s scoring capacity is not the limiting factor. Instead, these results define a clear physical boundary for the protocol: reweighting is highly valid provided the engineered mutations optimize or perturb the interface without completely re-routing the physical mechanism of dissociation.

Another fundamental source of error stems from the IFMs themselves, which are adept at extrapolating in structural space but are not guaranteed to work reliably in sequence space. This behavior is primarily dictated by their training paradigm. Models like ProteinMPNN take geometric features such as pairwise distances of backbone atoms as inputs, and added structural noise during training to capture stochastic local coordinate variations.^24^ However, because they are trained on scarce, individual PDB structures mapped to specific sequences, they cannot see the underlying evolutionary or homology context. Consequently, they can overlook sequence-space dependencies when evaluating engineered variants, suggesting that integrating a evolutionary prior from language model could help with this training data scarcity and significantly improve their prediction for design tasks. Despite this limitation, the correlation we observed between MAHLER and IfMetaD calculations is on par with thermodynamic predictors widely deployed in industry, establishing what is considered practically usable for prospective engineering workflows. For context, state-of-the-art molecular docking methods typically yield a Spearman *r* of approximately 0.6 when compared to experimental binding free energies^33^. Similarly, recent IFM-based binding free energy predictors^32^ generally report correlations between 0.5 and 0.7 depending on the system; thus, the alignment of our kinetic framework with these benchmarks underscores that we are likely reaching the current accuracy limit inherent to zero-shot IFMs.

To contextualize this dependency, an analogy can be drawn to the development of free energy perturbation (FEP) methods. While Zwanzig proposed the original theoretical framework for FEP^34^ in 1954, it did not become a practical tool until computational hardware advanced^11^ and force field development^35–37^ reached a critical threshold of accuracy. Today, relative FEP methods are reaching a practical 1-2 kcal/mol accuracy, and absolute FEP methods are catching up quickly^38, 39^. Just as FEP required highly calibrated physical force fields and faster computation to become mainstream, the MAHLER framework relies on the continued maturation of IFMs to serve as structural foundation models or “force fields”, and continued development in data-driven CV discovery.

In summary, the MAHLER framework establishes a scalable hybrid paradigm that successfully overcomes the throughput barriers historically associated with predicting macromolecular dissociation kinetics. By combining IFM-based trajectory reweighting with physical sampling, we achieve relative residence-time predictions with the necessary accuracy for high-throughput screening at a manageable computational cost. This breakthrough unlocks the ability to explore vast mutant libraries previously unimaginable with wet-lab experiments or explicit physics-based simulations alone. Consequently, our work significantly advances the scope of inverse-folding reweighting protocols^17^ and demonstrates that kinetics-aware antibody design is no longer just a theoretical goal, but a scalable computational reality. Ultimately, our results provide the practical framework needed to transition kinetic optimization from a niche challenge into a standard pillar of prospective immunotherapy design.

## Methods

### 1 Interface residue distances and interface fraction of native contacts

Interface residue distances (*d*^*IR*^) and interface fraction of native contacts *Q*-value can then be defined with the experimental structure. Native contacts were defined as any heavy atom pairs across the Ag-Ab interface that are closer than 5 Å. The fraction of native contacts was defined as in Ref. 40 to be a continuous function that maps a structure to a value between [0, 1].

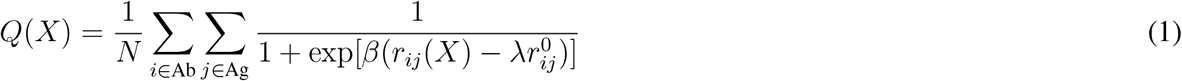

where *β* = 5 Å^−1^ and *λ* = 1.8. The reference distance 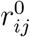 is taken from the experimental structure. Interface residues are defined as those residues with at least one atom involved in native contacts (Figure S5). The *d*^*IR*^ is calculated as the distance between the center of mass (COM) of C_*α*_ atoms of Ag interface residues and that of Ab interface residues.

### 2 Structural hypothesis generation with AlphaFold-Multimer

The AgAb complex structures were sampled with AF-M 2.3 as implemented in ColabFold 1.5.5, together with its MMseq2 server.^41, 42^ We used AF-M for its more commercially permissive license and easily tunable MSA depth to control diversity versus quality. The generated structures were input into the af2rave protocol to learn a reaction coordinate.

A common way to introduce diversity into AlphaFold predictions is to limit the MSA information available to the model^43^. However, we observed that when reduced MSA was used for AF-M, the quality of the structures was significantly compromised. Often, the individual chains were still well-folded, but we observed unphysical clashes between chains. This problem was less common in single chain predictions. Introducing templates significantly reduced structural diversity but improved the quality. Balancing all these factors, we applied a three-step strategy to maintain both diversity and quality.

First, we supplied a template, but for the antigen *only*. Antigens in resolved experimental structures often only contain fragments interacting with the antibody. Providing a template for the antigen significantly improved the quality of the folded antigen and reduced clashes. However, if a template for the antibody was also provided, diversity in loop structures and binding epitopes disappeared.

Second, we increased the number of recycles to five and saved all intermediate structures after each recycle. Each recycle of AF-M Evoformer will update the pair representation of the protein, sometimes changing the relative position of chains^44^. This strategy allowed us to extract more information from the inference process.

Third, we applied a steric clash filter to filter out structures with dangerously close heavy atoms (closer than 1.0 Å). This served as the last guard against unphysical structures.

For 2CMR, for example, our protocol eventually yielded 11450 structures from four MSA depths (8:16, 16:32, 32:64, and 512:5120) with 128 seeds for each of the five AF-M models. This setup performed 128 × 5(models) × 4(MSA depths) = 2560 inferences, but with intermediate structures saved for each recycle, we more than quadrupled the number of outputs to 11450. Finally, 4958 of them (43.3%) passed the steric clash filter. The system 3HFM yielded a similar 14919 structures, and only 1907 (12.8%) passed the steric clash filter.

### 3 Infrequent Metadynamics

Infrequent metadynamics (IfMetaD) builds upon the standard metadynamics (MetaD) framework to estimate residence times^12^. MetaD simulations drive systems out of local minima by depositing a repulsive, history-dependent bias potential, allowing rare events such as AgAb dissociation to be sampled within accessible simulation timescales^45, 46^. The core premise of IfMetaD is that if the bias deposition is slow enough to avoid corrupting the transition state region, the acceleration factor *α* (defined as the ratio of the physical residence time *τ* to the simulation time *τ*_*M*_) can be calculated with a simple relation:

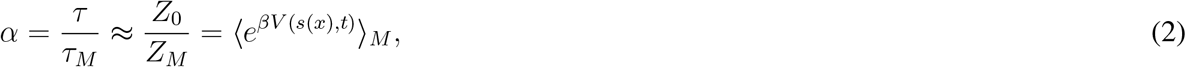

where *Z*_0_ and *Z*_*M*_ are the partition functions of the reactant in the unbiased and metadynamics ensembles, respectively, and *V* (*s*(*x*), *t*) is the time-dependent bias acting on the collective variables *s*(*x*).

In practice, the physical clock time is recovered by discrete sums over the metadynamics trajectory steps:

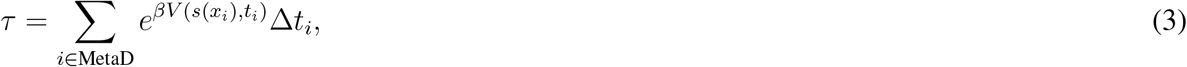

where {*x*_*i*_} represents trajectory frames saved at interval Δ*t*_*i*_.

Because AgAb dissociation is a stochastic process governed by Poisson kinetics, the residence times are random variables that follow an exponential distribution.^47^ Therefore, IfMetaD is performed using multiple independent replicas. The resulting distribution of dissociation times is then fit to an exponential cumulative distribution function (CDF) (Figure 1b) to yield a robust statistical estimate of *k*_off_.

In our calculations, we started from a structure equilibrated in the AlphaFold2-RAVE process, and used the same simulation protocol as described in the next section.

### 4 AlphaFold2-RAVE

Successful IfMetaD relies on a well-chosen set of collective variables (CVs). While the definition of an “ideal” CV varies depending on whether one aims to capture the reaction mechanism or the transition state^12, 48, 49^, IfMetaD imposes specific requirements based on Equation (2). This equation links the theoretical acceleration to the partition function of the reactant ensemble only^50, 51^. This implies that the primary contributor to the acceleration factor is the sampling within the reactant basin^15^, effectively removing the need for detailed knowledge of the transition state. Consequently, a satisfactory CV must facilitate thorough sampling of the bound AgAb complex. Variables with significant degeneracy, such as RMSD or center-of-mass distances, are therefore unsuitable as they fail to resolve the basin’s distinct metastable states (Supplementary Information section A.1).

Generated structures from the previous step were input into the feature selection module of the af2rave package^23^. The algorithm first selected the 200 interatomic distances with the highest coefficient of variation from the defined key atoms. Key atoms included C_*α*_ of the interface residues, but may extend to other atoms if prior knowledge is available about the specific system (Supplementary Information section A.2 and Ref. 23). Principal component analysis (PCA) reduced the dimensionality of these 200 distances to 20 components, after which *K*-medoids clustering identified 15 representative structures. Seven structures with *Q <* 0.85 were discarded. The remaining eight structures, plus the experimental crystal structure, were each simulated for 50 ns, yielding a cumulative sampling time of 0.45 µs. During these simulations, the 200 selected pairwise distances were recorded. The AMINO (Automatic Mutual Information Noise Omission) module^52^ then analyzed these time series to remove redundancy, narrowing the selection to ~ 20 distinct distances (Figure S5). Finally, the SPIB module (using Δ*t* = 0.1 ns and default parameters) processed these inputs to learn a 2D latent space. The resulting collective variables (CVs) were linear combinations of these interatomic distances^19^.

Metadynamics simulations were performed in this latent space using GROMACS 2023^53^ patched with PLUMED 2.9^54^. A Gaussian bias with a height of 0.3 kcal/mol and a width of 0.05 was deposited every 10 ps on the two latent variables, with a bias factor of *γ* = 15. Simulations were monitored via *d*^*IR*^ and *Q* (though not biased) and were terminated when the dissociation criteria (*d*^*IR*^ greater than initial value plus 3 Åand *Q <* 0.5) were both met. The acceleration factor from Eq. 2 was calculated on the fly by PLUMED. The final dissociation time was computed as the product of the acceleration factor and the simulation time. Trajectories were saved to disk every 10 ps.

The resulting two CVs encode essential information about the dissociation process through extrapolation. Crucially, the latent space was learned using MD data from bound structures only (*Q >* 0.85). To test how well this space extrapolates to the dissociation process, we ran 30 exploratory well-tempered metadynamics (WTMetaD) simulations by aggressively biasing the interface residue distance (*d*^*IR*^) and *Q* (defined in Methods). While these *ad hoc* CVs are inappropriate for statistical sampling (Supplementary Information section A.1), they quickly drive the system to dissociate. When these dissociation trajectories were projected onto our learned latent space, the two latent variables correlated smoothly with the *Q*-value and *d*^*IR*^ (Figure S4) of the sampled structures.

### 5 Reweighting and off-rate calculation

The derivation of the IFM reweighting framework begins with Bayes’ theorem, which allows us to relate the conditional probabilities of a wild-type (WT) sequence *S* and a mutant sequence *S*′ for a particular conformation *x*:

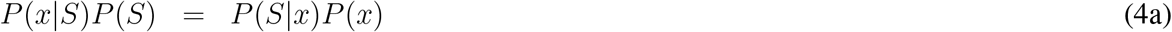

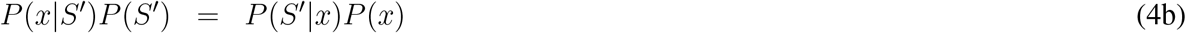

Dividing Equation (4a) by Equation (4b) and rearranging yields the master reweighting relationship:

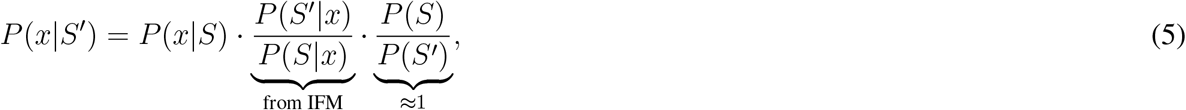

Consequently, sampling from the target mutant ensemble *X*′ ~ *P* (*x*|*S*′) can be achieved by reweighting samples from the reference WT distribution *X* ~ *P* (*x*|*S*), for which simulations have already been performed.

This theoretical basis for the reweighting was first derived by Brotzakis *et al*. in Ref. 17. However, our protocol departs from their approach in two key respects. First, in their work, they biased the RMSD of the whole system compared to the experimental structure as the only CV. As described before and in Supplementary Information section A.1, this CV does not help distinguish different metastable states and is not the best choice of CV when sampling dissociation. In our work, we biased a machine-learned two-dimensional CV from af2rave for better sampling.

Second, we approximate the ratio of sequence priors *P* (*S*)*/P* (*S*′) as unity^32^, instead of using a protein language model. We justify this approximation with two main arguments. First, while previous studies have modeled this term using protein language models (pLMs)^17, 32^, pLMs are trained on natural evolutionary databases and may not accurately reflect the statistical trends of artificial protein design, where sequences often deviate significantly from evolutionary manifolds^55^. From a strictly statistical perspective, in the absence of evolutionary constraints, there is no *a priori* reason to favor one sequence over another. Second, recent work by Frellsen *et al*.^32^ analyzing the thermodynamic interpretation of inverse folding models found minimal difference in correlation with experimental data whether the prior term was included or not. Given these factors, we opted for the parsimonious approach of assuming a uniform prior, thereby avoiding unnecessary model complexity.

The reweighting factor in Equation (5) relies on the likelihood of a sequence conditioned on a specific structure, *P* (*S*|*x*), which is directly estimated using scores from an IFM, where we adopted ProteinMPNN^24^. In practice, each trajectory of the WT simulation is independently reweighted with the mutant sequence to predict a new dissociation time. Combining this structural weight with the expression of the acceleration factor in Eq. 2, we get the expression for reweighted time:

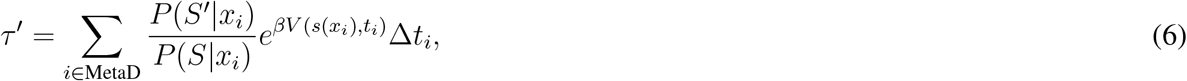

where *i* iterates over all saved MetaD trajectories.

We employed ProteinMPNN^24^ to estimate the negative log-likelihood − log *P* (*S*|*x*). We used the pre-trained model with a context length of 48 amino acids and a training noise level of 0.2 (`v 48 020`). In this context, *S* corresponds to the wild-type (WT) sequence, and *S*′ corresponds to the specific point mutant under study.

Once dissociation times were obtained for each replica, either directly from IfMetaD or via reweighting, we constructed the empirical CDF of the dissociation events over time (Figure 1b). This distribution was fit to a theoretical exponential CDF, *P* (*t*) = 1 − *e*^−*t/τ*^, to extract the characteristic residence time *τ*. The dissociation rate was then calculated as *k*_off_ = 1*/τ*.

The error bars in all main text figures represent the standard deviation obtained via bootstrapping. We performed resampling with replacement on the dissociation events to generate 1000 bootstrap samples (each containing 300 events). A new CDF was computed and fit for each sample to determine the standard deviation of the resulting *τ* distribution. The error bars in Figure 3b correspond to the standard error of the mean across the three independent MD replicas.

## Code Availability

An open source package for our protocol is available at github.com/tiwarylab/mahler. The data from the IfMetaD was deposited at doi.org/10.5281/zenodo.17886490 and a demonstration notebook of the reweighting code is available at https://colab.research.google.com/github/tiwarylab/mahler/blob/main/notebook/MAHLER_Reweighting.ipynb. This notebook downloads the trajectory and structure from Zenodo and calls the MAHLER package to reweight it. It takes around 50 minutes to run on a NVIDIA A100 runtime and reproduces Figure 2f in the main text.

## Acknowledgements

P.T. was supported by the National Institute of General Medical Sciences of the National Institutes of Health under Award Number R35GM142719. D.T. was supported by SandboxAQ. P.T. is an investigator at the University of Maryland-Institute for Health Computing, which is supported by funding from Montgomery County, Maryland, and The University of Maryland Strategic Partnership: MPowering the State, a formal collaboration between the University of Maryland, College Park, and the University of Maryland, Baltimore. We thank UMD HPC Zaratan and NSF ACCESS (project CHE180027P) for computational resources. The authors thank Akashnathan Aranganathan, Dr. Shams Mehdi, Suemin Lee, Dr. Xinyu Gu, and Dr. Yunrui Qiu for helpful discussions.

## Supplementary Information

### A Supplementary Discussions

#### A.1 On selection of CVs and exploratory MetaD

As noted in the Methods section, we noted that collective variables (CVs) such as the *Q*-value or AgAb distance are often inappropriate for rigorous enhanced sampling. While the definition of an “ideal” CV depends on specific sampling or analysis goals, a fundamental requirement is the ability to resolve distinct metastable states^56^.

By this metric, geometric descriptors like RMSD, antigen-antibody distance, or the fraction of native contacts are often suboptimal. These variables typically suffer from high and heterogeneous degeneracy; multiple distinct metastable states can map to the exact same scalar value (e.g., the same RMSD or distance). Consequently, these CVs frequently fail to capture the system’s slowest modes, leading to hysteresis and poor convergence.

In this work, we employed the interface residue distance and interface *Q*-value strictly to drive the system toward dissociation, not to recover a converged free energy surface, or for kinetics calculation. These variables do not guarantee that the system visits all relevant intermediate states.

This limitation is evident in our 30 replicas of exploratory MetaD (Figure S1), where the bias potentials deposited to induce dissociation varied significantly in both height and shape across replicas. Such variance indicates insufficient sampling of orthogonal degrees of freedom. For reference, these simulations used a Gaussian bias with a height of 0.3 kcal/mol, widths of 0.2 Å(distance) and 0.02 (*Q*-value), and a bias factor of 15.

#### A.2 On the choice of key atoms

The feature selection algorithm is detailed in Ref. 23; here, we summarize its specific application to this system.

We define a set of key atoms comprising *N*_*g*_ atoms from the antigen and *N*_*b*_ atoms from the antibody, generating a total of *N*_*g*_*N*_*b*_ pairwise distances. To identify features that capture the conformational diversity predicted by AF-M, we rank these distances by their coefficient of variation (standard deviation divided by the mean) across the generated ensemble. The top *s* features (here, *s* = 200) are used to cluster the conformations. From this reduced pool, the AMINO algorithm identifies a sparse, non-redundant subset (here, 19 distances) to learn the latent space.

For the candidate pool, we selected the C_*α*_ atoms of all interface residues (as defined in Methods). We restricted the selection to surface atoms for two physical reasons. First, the antibody core is generally rigid, so biasing long-range distances involving core atoms risks forcing unphysical deformations rather than valid conformational changes. Second, distances involving atoms away from the interface are typically much longer than those between interface pairs. Geometrically, when the radial degree of freedom is fixed, a larger radius results in a larger spherical shell. This increased angular freedom introduces higher degeneracy, rendering long-range distances less informative for distinguishing specific metastable states.

### B Supplementary Methods

#### B.1 Structures generated with AlphaFold-Multimer

We projected the structures generated by AF-M by several metrics. A few conclusions can be drawn from these results.

First, across the three accuracy predictors provided by AF-M, none of them is decisive in determining the correctness of the structure (Fig. S2), characterized by a high interface *Q*-value. Even ipTM scores are noisy in this sense. RMSD to experimental structures as a metric often ignores the existence of alternative binding epitopes.

Second, the dependency of structural quality, characterized by steric clashes, is complicated in terms of MSA depth (Fig. S3).

Different MSA depths give very different patterns. For example, full MSA tends to give higher quality structures in all three metrics. Shallower MSA usually gives a longer tail in these figures, but nearly none of them pass the steric clash filter. It shows that there is probably not a single “best” MSA depth to gain diversity.

Despite this longer tail, it doesn’t mean shallower MSA necessarily is better at discovering alternative modes, as demonstrated in the case of ADK in Ref. 23. For example, two “islands” of structures that are around 8 Å and 11–16 Å away from experimental structures appeared only in full MSA predictions. However, none of them passed the steric clash filter as well, indicating a clashing binding epitope.

#### B.2 Extrapolation of SPIB latent space

Here in Figure S4, we report the projection of the 30 exploratory MetaD trajectories from 2CMR in the latent space, colored by fraction of native interface contacts. Here we juxtaposed the same projection colored by interface residue distances (left) and the and fraction of native contacts (right).

Our results show that both in terms of interface residue distances (*d*^*IR*^) and *Q*-value, our latent space projects well, as a supplement to Fig. 1c.

#### B.3 Mean score calculation

Figure 3b showed the mean score from three unbiased MD runs. Trajectories were saved every 0.1 ns and scored with ProteinMPNN. The relative score {*s*_*i*_} from ProteinMPNN is a log-likelihood.

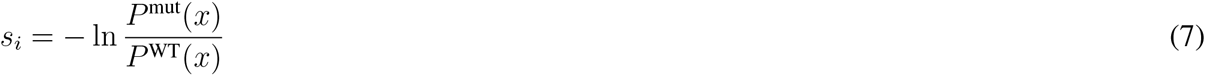

When setting *V* = 0 and that *e*^*βV*^ = 1 in Equation (6), a meaningful mean value can be calculated as

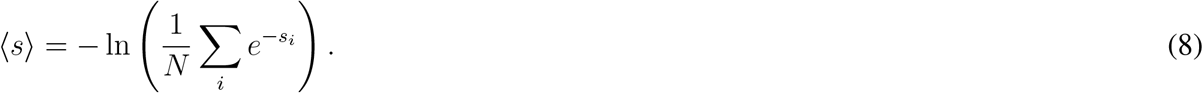

**Figure S1.**
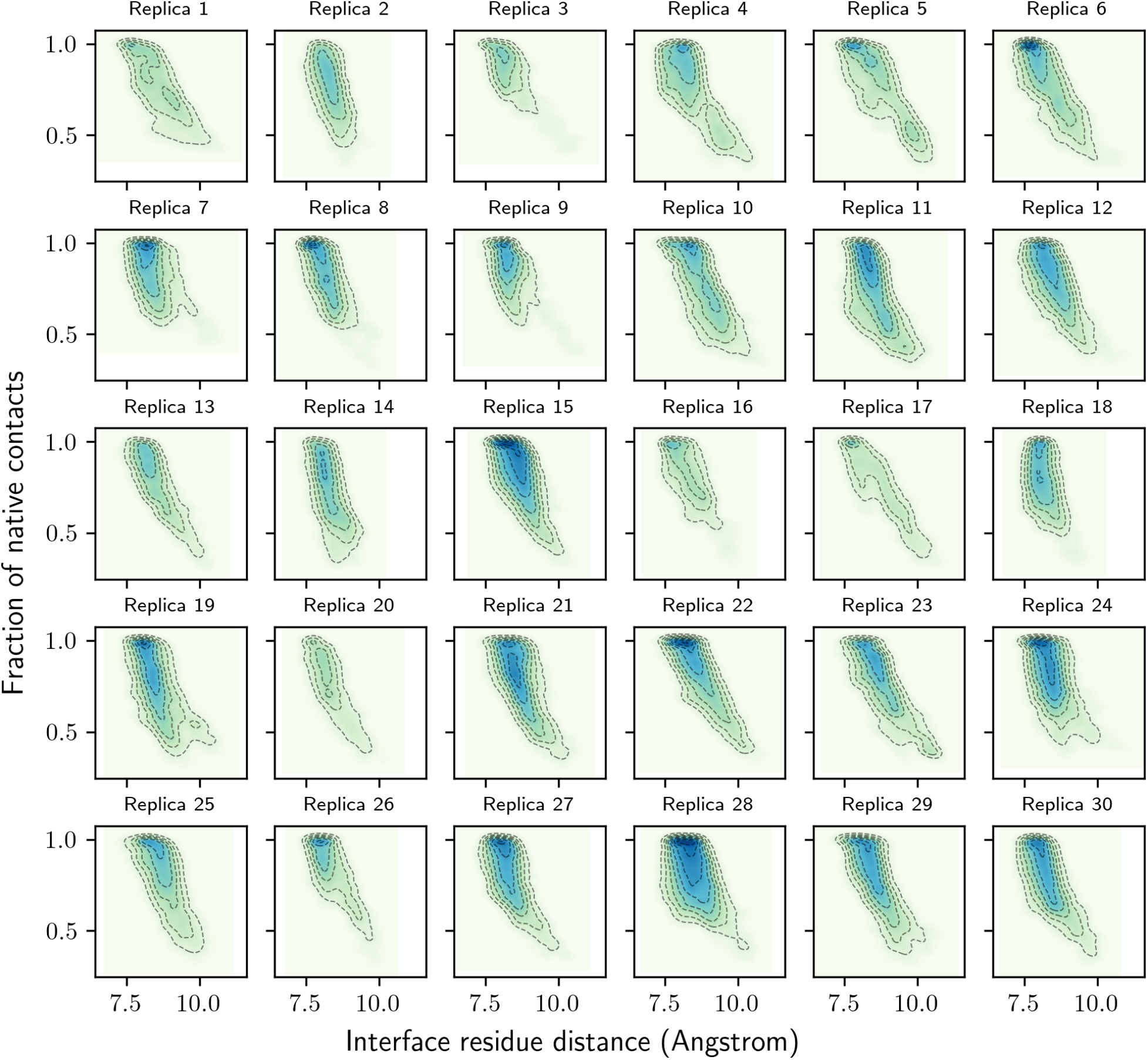
Hills deposited for the exploratory MetaD. 30 replicas of exploratory MetaD were run for 2CMR and were projected onto the latent space. Contour lines are spaced 1.0 kcal/mol away from each other. Color intensity corresponds to the cumulative density of deposited bias

**Figure S2.**
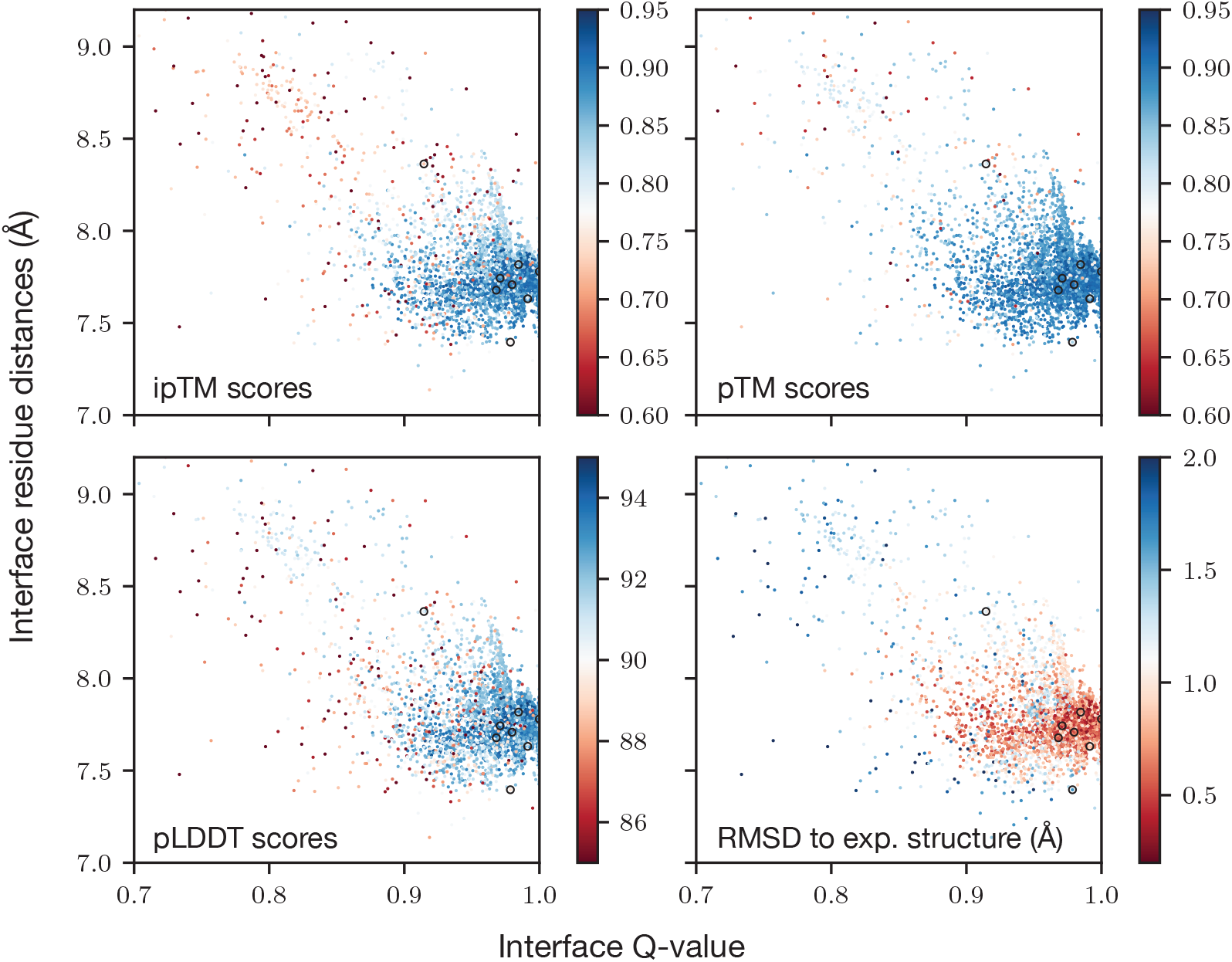
Qualities of AF-M generated 2CMR structures. AF-M can generate slightly dissociated structures, bypassing the need for additional sampling. All structures that passed the steric clash filter were projected by the interface *Q*-value and interface residue distances. Selected representative structures are shown in black circles. Three AF-M predicted metrics (ipTM, pTM, and pLDDT) were used to color the points. Black circles show the selected seed structures for MD.

**Figure S3.**
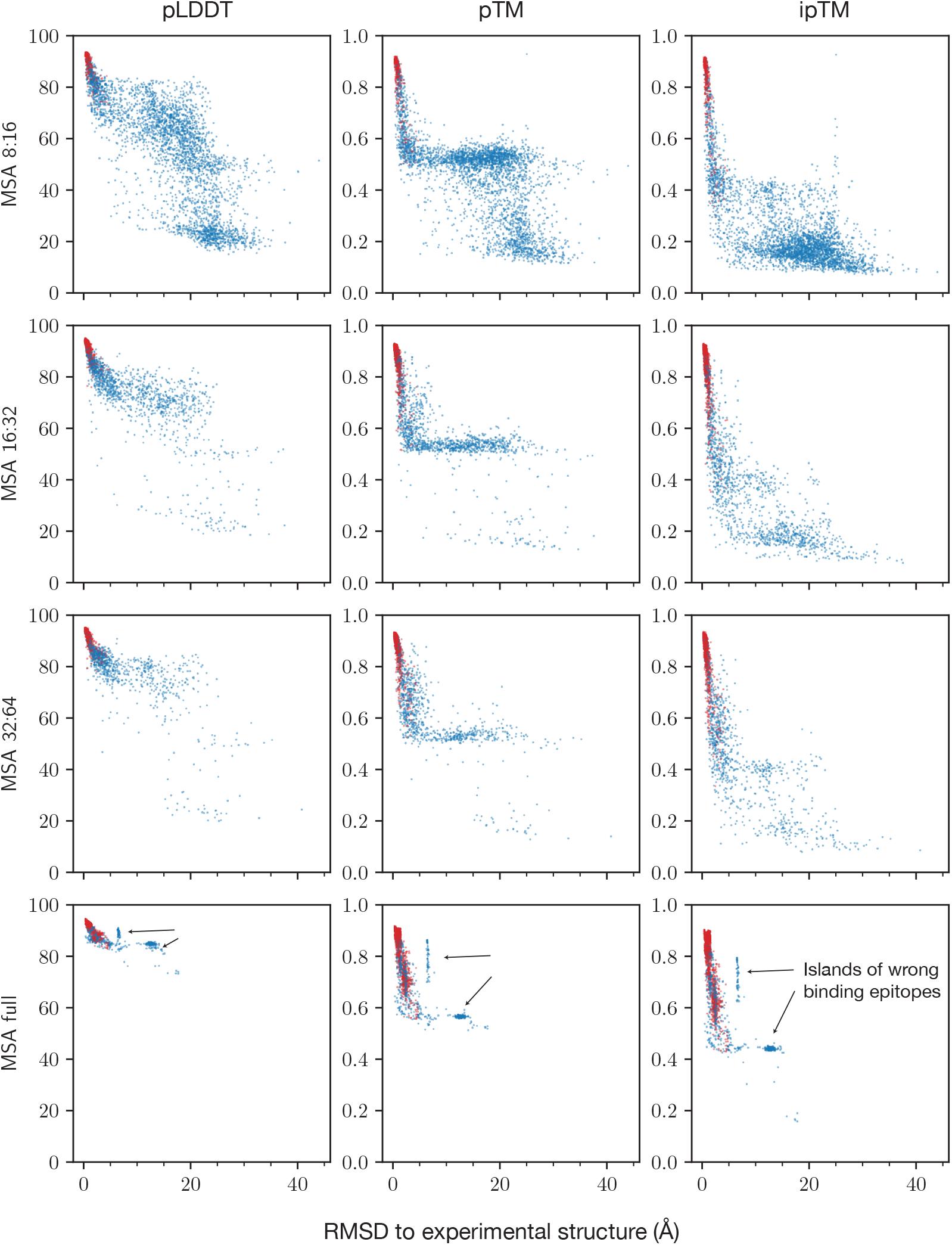
Effect of MSA depth and steric clash filter. This figure provides a profile of the generated structures. The red points are those passing the steric clash filter at 1 Å.

**Figure S4.**
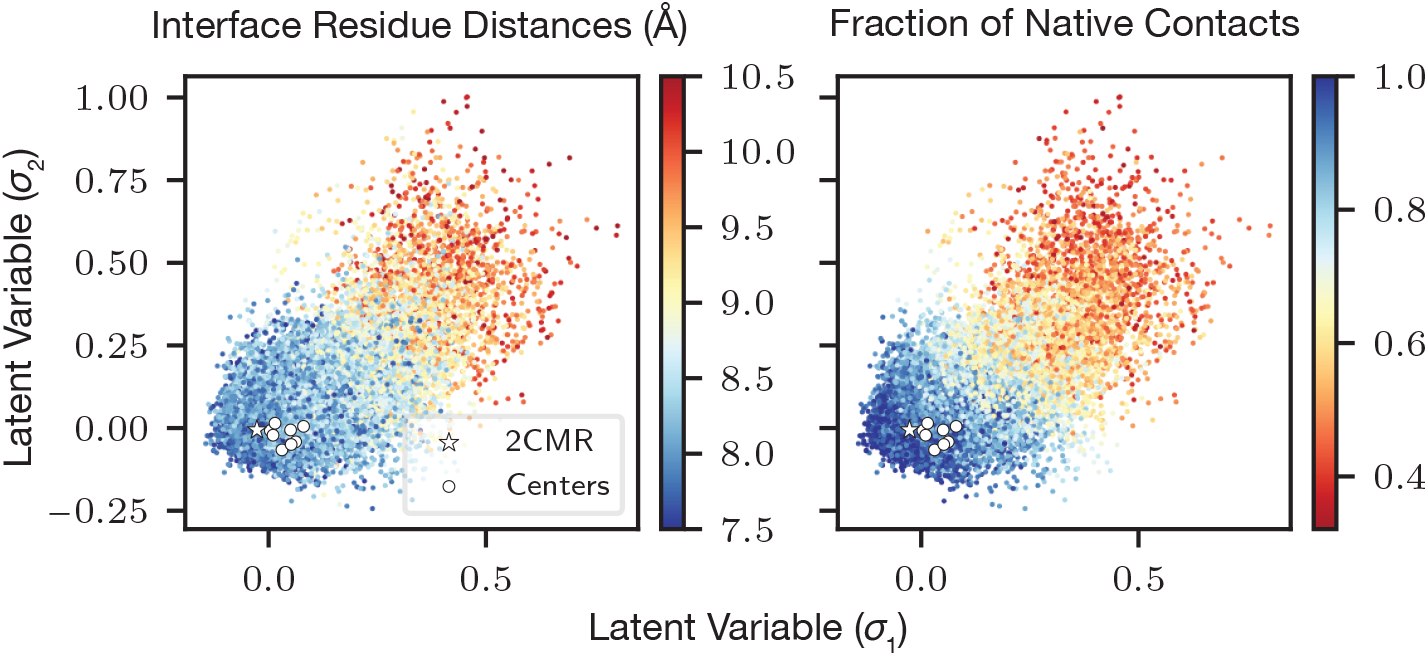
Projection of exploratory 2CMR MetaD trajectories on SPIB latent space.

**Figure S5.**
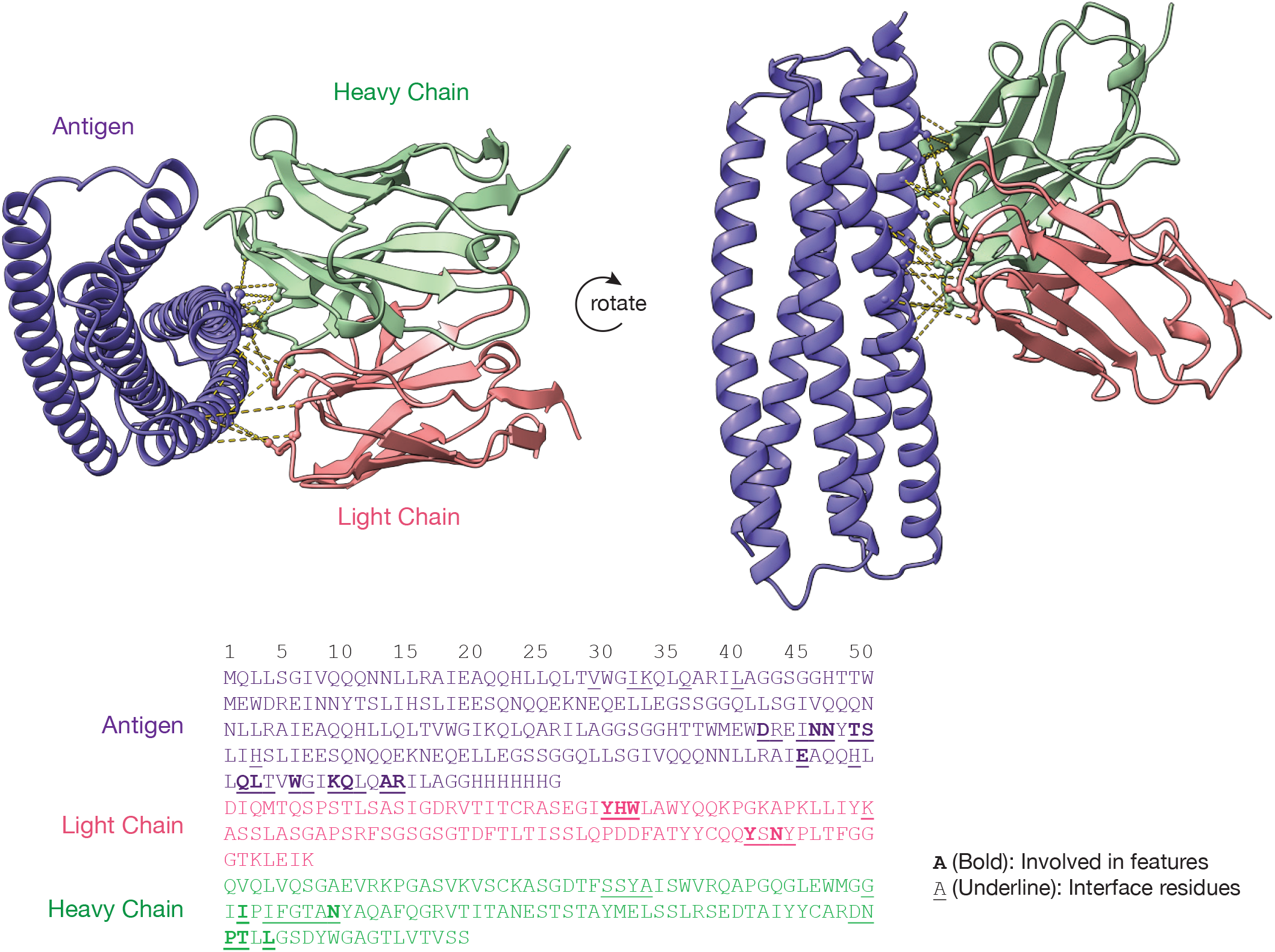
Sequence and interface of the 2CMR system. Upper: visualization of the 19 distances used as features. Lower: sequence for the system and key residues involved.

